# Group A Streptococcal Collagen-like Protein 1 Restricts Tumor Growth in Murine Pancreatic Adenocarcinoma and Inhibits Cancer-Promoting Neutrophil Extracellular Traps

**DOI:** 10.1101/2024.01.17.576060

**Authors:** Emily A. Henderson, Abby Ivey, Soo Choi, Stell Santiago, Dudley McNitt, Tracy W. Liu, Slawomir Lukomski, Brian A. Boone

## Abstract

Pancreatic ductal adenocarcinoma (PDAC) is a lethal cancer associated with an immunosuppressive environment. Neutrophil extracellular traps (NETs) were initially described in the context of infection but have more recently been implicated in contributing to the tolerogenic immune response in PDAC. Thus, NETs are an attractive target for new therapeutic strategies. Group A Streptococcus (GAS) has developed defensive strategies to inhibit NETs. In the present work, we propose utilizing intra-tumoral GAS injection to stimulate anti-tumor activity by inhibiting cancer-promoting NETs. Injection of three different M-type GAS strains reduced subcutaneous pancreatic tumor volume compared to control in two different murine PDAC models. Limitation of tumor growth was dependent on streptococcal collagen-like protein 1 (Scl1), as isogenic mutant strain devoid of Scl1 did not reduce tumor size. We further show that Scl1 plays a role in localizing GAS to the tumor site, thereby limiting the systemic spread of bacteria and off-target effects. While mice did elicit a humoral immune response to GAS antigens, tested sera were negative toward Scl1 antigen following intra-tumoral treatment with Scl1-expressing GAS. M1 GAS inhibited NET formation when co-cultured with neutrophils while Scl1-devoid mutant strain did not. Recombinant Scl1 protein inhibited NETs *ex vivo* in a dose-dependent manner by suppressing myeloperoxidase activity. Altogether, we demonstrate that intra-tumoral GAS injections reduce PDAC growth, which is facilitated by Scl1, in part through inhibition of cancer promoting NETs. This work offers a novel strategy by which NETs can be targeted through Scl1 protein and potentiates its use as a cancer therapeutic.

## 1 Introduction

Pancreatic adenocarcinoma (PDAC) is one of the most lethal cancers with a five-year survival rate of only 12% [1,2]. Late diagnosis and resistance to current treatment modalities contribute to this poor survival [3,4]. PDAC is an immunologically “cold” tumor, with minimal response to checkpoint immunotherapy [5, 6]. Therefore, there is a growing need for the development of strategies to induce an anti-tumor immune response and generate novel cancer therapies in this deadly disease.

William Coley, a bone osteosarcoma surgeon influenced by the work of German physician Wilhelm Busch, performed the first systematic study using streptococcal organisms as a cancer treatment strategy in 1891 leading to the development of Coley’s toxin, a concoction of heat-killed *Streptococcus pyogenes* (group A *Streptococcus*) and *Serratia marcescens*. Though his work fell out of favor with implementation of radiation and chemotherapy, evidence suggests Coley’s intuition of stimulating the immune system to elicit an anti-cancer response is effective [7]. Bacteria harbor a variety of structures and mechanisms that make them attractive candidates as therapeutic agents with some evidence of preclinical efficacy in PDAC [8]. Evidence suggests that group A *Streptococcus* (GAS) peptides [9] and superantigens [10] can elicit tumor regression. GAS therapy showed promise in instances of bladder cancer [11], though was less successful in preclinical trial of other cancers [12, 13]. Still, GAS utilizes several structures to evade innate immunity which offers mechanisms by which GAS can stimulate an anti-tumor response. One mechanism is the inhibition of neutrophil extracellular trap (NET) release [14]. NETs were initially described as a method for neutrophils to trap and kill bacteria and other infectious agents [15], however NETs are also upregulated in sterile inflammatory diseases including PDAC [16, 17]. Recent evidence suggests the significance of NETs in contributing to PDAC tumor growth, progression, metastasis, and promotion of an immunosuppressive tumor environment [18, 19, 20, 21]. Moreover, NET levels are a prognostic indicator of PDAC patient survival and treatment outcomes [22, 23]. GAS has the capacity to evade NET degradation through the activity of streptococcal collagen-like 1 protein which prevents NET release [14, 24], though the use of GAS to limit NETs has yet to be explored in the context of cancer.

Herein, we demonstrate GAS as a potential therapeutic agent in two murine models of PDAC. We also propose the therapeutic significance of streptococcal collagen-like-1 (Scl1) protein as an essential factor in GAS anti-tumor response that supports tumor colonization, and acts as a potent inhibitor of NET activity through reduction of myeloperoxidase activity. These findings have significant implications in the exploration of recombinant Scl1 as a novel therapy to limit PDAC progression.

## 2 Materials and Methods

### 2.1 Animals

Age and gender matched C57BL/6 wild-type (WT) mice (8-weeks old) were purchased from the Jackson Laboratory (Bar Harbor, ME, USA). All animal experiments were approved by the Institutional Animal Care and Use Committee of West Virginia University, protocol number 1602000144_R1.5 and 180918204 and complied with the use of experimental animals published by the National Institutes of Health.

### 2.2 Cell lines and cell-culture

Panc02 cells derived from a pancreatic adenocarcinoma model in male C57BL/6 mice were obtained from the National Cancer Institute. KPCY6422c1 cells derived from mouse pancreatic neoplasms were purchased from Kerafast (Kerafast, MA, USA). Panc02 cells were cultured in RPMI 1640 media supplemented with 10% fetal bovine serum and 1% Penicillin-Streptomycin (Hyclone, Logan, UT, USA). KPCY6422c1 cells were cultured with Dulbecco’s Modified Eagle Medium with 10% fetal bovine serum and 1% glutamine. Cells were grown at 37°C in a humidified 5% CO_2_ atmosphere.

### 2.3 Murine subcutaneous tumor model

Age and gender-matched 8-week-old C57BL/6 WT mice were injected with 5×10^5^ Panc02 or KPCY6422c1 cells into the right flank. Once tumors reached 7 mm (10-12 days post tumor implantation), tumors were injected with group A *Streptococcus* as outlined below or PBS. Caliper measurements of tumors were performed every other day until experimental endpoint. Length (L) and width (W) of tumor were measured and tumor volume in mm^3^ was calculated using equation for spherical ellipsoid (V_T_=0.5*L*W^2^) [25]. Tumors and spleens were harvested, weighed, homogenized, and cultured on blood agar (**Fig. S1**).

### 2.4 Generation of recombinant proteins

Recombinant rScl1 proteins were generated as described previously [26, 27, 28]. Briefly, rScl1-encoding clones were generated in the *Strep*-tag II cloning, expression, and purification system using the *E. coli* vector pASK-IBA2 designed for periplasmic expression (IBA-GmbH, Geottingen, Germany); rScl1.1 was derived from the Scl1 protein in M1-type strain, rScl1.28 in M28-type strain, and rScl1.41 in M41-type strain. Clones were verified by sequencing and rScl1 polypeptides by N-terminal sequencing. Bacterial cultures were grown in Luria-Bertani medium (LB broth, Miller) and protein expression was induced with anhydrotetracycline (0.2 μg/mL) for three hours. The periplasmic fraction was collected following treatment with CellLytic B Cell Lysis reagent (Sigma, B7435) and purified on *Strep*-Tactin Sepharose.

rEDA was produced intracellularly using the pQE-30 His tag purification system (Qiagen) in *E. coli* JM-109 [29]. *E. coli* was cultured in LB broth supplemented with ampicillin (100 μg/mL) and protein expression was induced with 1 mM isopropyl β-D-thiogalactopyranoside for three hours. Cells were harvested and lysed in lysis buffer (50 mM Tris/HCL, pH 8.0, 50 mM NaCl, 2 mM MgCl2, 2% Triton X-100, 10 mM beta-mercaptoethanol, 2 mg lysozyme, 1 mM EDTA-free protease inhibitor cocktail tablet, 1 mM phenylmethylsulfonyl fluoride (PMSF), 100U DNaseI). Supernatant was collected following centrifugation, and rEDA was purified using nickel-nitrilotriacetic acid-agarose resin (Qiagen).

Following purification, rScl1 proteins and rEDA were analyzed by SDS-PAGE (**Fig. S2**). Proteins were dialyzed in 25 mM HEPES pH 8.0 and stored at -20°C.

### 2.5 Group A Streptococcus strain preparation and injection

The M1-type GAS strain MGAS5005 Δ*scl1* mutant was generated by allelic replacement mutagenesis as described previously [30]. Briefly, *scl1.1* allele was cloned into an *E. coli* vector and coding sequence was replaced with a nonpolar spectinomycin resistance cassette spc2, resulting in pSL134 plasmid suicide in GAS. MGAS5005 cells were electroporated with pSL134 and spectinomycin-resistant colonies were screened by PCR for a single amplification product indicating double cross-over event; mutant candidates were then verified by sequencing and loss of Scl1 protein was confirmed by western immunoblotting with specific anti-Scl1.1 Abs.

Complementation of M1 was performed as previously described [26] using GAS-*E. coli* shuttle plasmid. The DNA fragment expressing *scl1.1* was PCR-amplified from genomic DNA of MGAS5005 and cloned into pSB027 generating plasmid pSL620. pSL620 was introduced into MGAS5005 Δ*scl1.1* mutant by electroporation and selected on Brain Heart Infusion (BHI) agar with 10 μg/mL chloramphenicol; Scl1.1 expression was confirmed by western immunoblotting as above.

M1 wild type (WT), Δ*scl1.1* isogenic mutant, Δ*scl1.1::620* complemented mutant, M41 strain MGAS6183, and M28 strain MGAS6143 group A *Streptococcus* were grown overnight at 37LC with 5% CO_2_ on BHI agar solid media supplemented with antibiotics, as needed. Plated bacteria were inoculated into Todd-Hewitt Yeast (THY) broth and incubated until OD_600_ reached ∼0.5. Following centrifugation, bacterial cells were washed once with PBS, centrifuged again, and diluted in PBS to designated concentrations (M1; 1×10^5^, 1×10^6^, 1×10^7^; M41 and M28; 1×10^4^, 1×10^5^, 1×10^6^). Indicated inocula of bacteria in 0.1-mL volume were injected into subcutaneous tumors established in C57BL/6 mice. Control mice received intra-tumoral PBS injections at equal volume.

### 2.6 ELISA and western blot analysis

To assess mouse seroconversion to streptococcal antigens, blood was collected from tumor burdened mice 14 days following GAS injection or PBS controls. For whole-cell ELISA, M1 wild-type bacteria harvested at OD_600_ ∼0.5 and washed as above in PBS. Cells were finally suspended in bicarbonate buffer, pH 9.6; for coating, 1×10^7^ CFU were added into wells of 96-well high binding microplates (Thermofisher, 15041) and incubated at 4LC overnight [31]. Wells were washed with tris-buffered saline (TBS)-1% BSA and blocked in this reagent at 37LC for 2 hours. Primary mouse sera were added at a 1:100 dilution in TBS-1% BSA. Anti-M1 and anti-Scl1.1 rabbit antibodies were used as positive controls. Plates were incubated at 37LC for 2 hours. Anti-mouse or anti-rabbit HRP-conjugated secondary Abs at 1:1000 dilution were added and incubated at room temperature for 1 hour; then, wells were washed and ABST substrate was added and immunoreactivity was read at 415 nm.

Western immunoblotting was carried out against GAS cell wall-associated protein fraction prepared from MGAS5005 and separated on 15% SDS–PAGE, as described [30]. Log-phase GAS cells were harvested and resuspended in 20% sucrose–10 mM Tris (pH 8.0), 1mM ethylene diamine tetra-acetic acid buffer containing 5 U of mutanolysin,100 μg of lysozyme, and 100 mM PMSF. After digestion at 37°C for 1 h and centrifugation, the supernatants were used for subsequent analyses. The primary antibodies were the same sera used above in whole cell ELISA prepared from mice bled at the endpoint of experiments. The control anti-GAS-antigen antibodies include: (i) anti-M1 protein rabbit Ab (1:2500 dil.) [32] and anti-Scl1.1 affinity purified rabbit Ab (1:1000 dil.) [30]. Secondary Abs used were, accordingly, goat anti-mouse IgG(H+L)- or goat anti-rabbit IgG(H+L)-HRP conjugated (Bio-Rad, 1721019), followed by chemiluminescent detection with Clarity Western ECL Substrates (Bio-Rad, 1705060).

### 2.7 Neutrophil extracellular trap assay and quantification

Bone marrow neutrophils were isolated from the femur of euthanized C57BL/6 mice as described previously [16, 33]. Bone marrow was collected, and cells were washed with RPMI-10 with 10% FBS and 1% PenStrep. Cells were separated using a density gradient centrifugation with histopaque 1077 and 1099. The layer between 1077 and 1099 was collected and plated onto a 96-well plate at 1.5×10^4^ cells per well in Hank’s Balanced Salt Solution. Neutrophils were incubated for thirty minutes at 37LC to allow for cell attachment. Recombinant Scl1 protein, HEPES, sterile PBS, or whole-cell bacteria were added at the designated concentration and incubated for 5-10 minutes. 50μM platelet activating factor (511075, Sigma Aldrich) was added for activation for 3 hours. Cells were fixed with 4% paraformaldehyde and stained for DNA with Hoechst reagent. NETs were visualized using an EVOS microscope at 10-40x objective. NETs were quantified by collecting the supernatant and measuring cellular free DNA using Quant-iT Picogreen (Invitrogen, Grand Island, NY) according to manufacturer’s protocol. 500,000 U/mL polymyxin B sulfate was added to the recombinant protein for all *in vitro* experiments to account for LPS contamination.

### 2.8 Myeloperoxidase assays and quantification

Recombinant Scl1 proteins were examined for MPO activity using the Myeloperoxidase Inhibitor Screening Assay Kit (700170, Cayman Chemical, Ann Arbor, MI). The proteins were reconstituted in PBS and used in the assay according to the manufacturer’s protocol.

MPO activity was also assessed by luminol bioluminescence imaging of isolated bone marrow neutrophils as described previously [34, 35]. Briefly, 1×10^5^ isolated neutrophils were added to a 96-well black walled plate and stained with 100mM luminol (Sigma-Aldrich, MO, USA) followed by stimulation with 500nM phorbol 12-myristate 13-acetate, 99+% (Thermo Scientific Chemicals). Immediately following stimulation, cells were imaged for 60 minutes using the Kino imaging system (Spectral Instruments Imaging, AZ, USA) at 37°C under 5% CO_2_ flow for 12 total acquisitions. Data was analyzed by ROI measurements using Aura software (Spectral Instruments Imaging, AZ, USA) imported into Excel (Microsoft Corp., WA, USA). Data was represented as total flux (photons/second) by averaging triplicate wells and calculating the area under the curve from images taken at 5-60 min.

### 2.9 Statistical analysis

All statistical analysis was performed on GraphPad Prism 10.1.0 (316). Experiments were analyzed for significance using one-way ANOVA with or without repeated measures with multiple comparison post hoc tests as indicated in figure legends. Significance is designated as *=p<0.05, **=p<0.01, ***=p<0.001, ****=p<0.0001.

## 3 Results

### 3.1 GAS limits subcutaneous Panc02 murine PDAC tumor growth

*S. pyogenes* serotype M49 has been shown to elicit anti-tumor growth activity *in vivo* against Panc02 tumors [36, 37]. We first sought to substantiate these findings and investigate whether this effect was observable in other M-types or was restricted to M49. We utilized three different strains of *S. pyogenes* (M1, M28, and M41) at three different inocula injected into subcutaneous murine PDAC tumors. One week following injection of 5×105 Panc02 cells into the right flank, male and female mice were randomized to intra-tumoral injection with low, medium, and high inocula of each GAS M type or PBS control at equal volume. Tumor burden was measured on day eight post-infection.

Intra-tumoral injection with all three strains exhibited minimal skin pathology on day 8 (**Fig. 1A**), while eliciting significant and comparable reductions in tumor volume and weight using all three strains compared to the PBS-treated tumors **(Fig. 1B, 1C).** Tumor reduction was largely inoculum-size dependent, except for M28, which was most efficacious using the medium inocula. Only two mice inoculated with the high M1 inoculum died from systemic spread of bacteria indicated by the growth of β-hemolytic GAS colonies on blood agar inoculated with blood and spleen homogenates from these mice (**Fig. S1**). Overall, we established that GAS of various M-types are capable to reduce Panc02-tumor growth *in vivo*.

**Figure 1.**
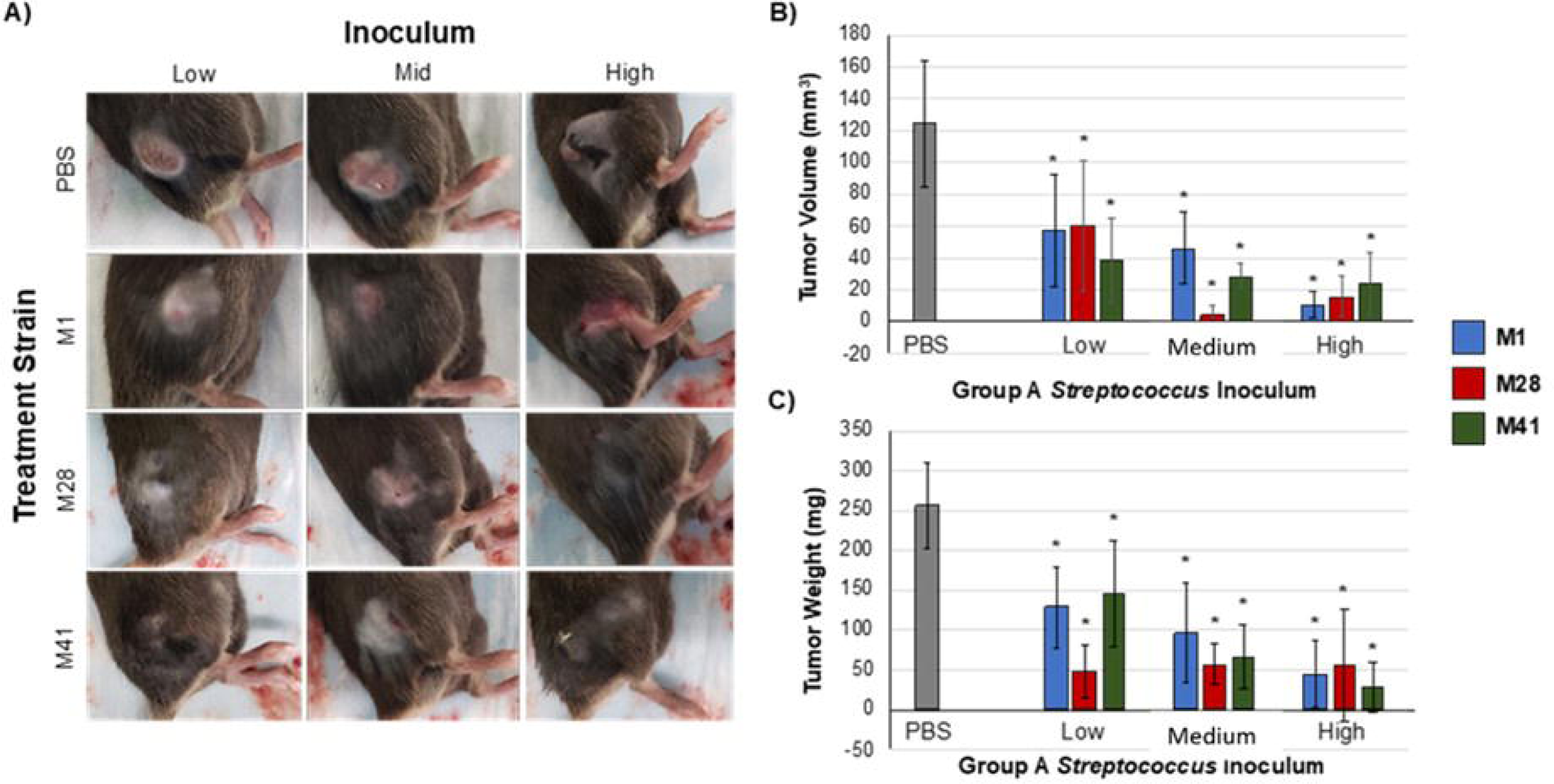
GAS injection limits the growth of subcutaneous murine Panc02 tumors. (A) Representative images of tumors and skin pathology associated with GAS injection. Pre-formed Panc02 tumors were injected intratumorally with GAS strains at low, medium, and high inocula (M1: 10^5^, 10^6^, 10^7^ CFU or M41/M28: 10^6^, 10^7^, 10^8^ CFU) or PBS at equal volume. Data shown were recorded on day 8 post-injection. (B) Comparison of tumor volume between experimental groups. Tumor dimensions were measured with a caliper on day 8 post-injection of GAS compared to PBS control. (C) Comparison of tumor weight between the groups. Tumors were collected and weighed on day 8 post-injection of GAS or PBS. *=p<0.05. n=5/group.

### 3.2 GAS injection limits the growth of subcutaneous KPC murine PDAC tumors

The *KRAS* gene is an oncogene that encodes the protein KRAS involved in regulating cell proliferation and survival. Mutations in *KRAS* can disrupt the switch between active and inactive states thus promoting cancer development [38]. Virtually all PDAC tumors harbor *KRAS* mutations and a majority also have mutations in P53, a tumor suppressor protein [3, 39]. Therefore, we investigated whether GAS M1 strain had similar anti-tumor effects using KPCY cells, a PDAC tumor lineage that mirrors the genetics of human PDAC [40]. We injected 10^6^ CFU, the medium-size inoculum in Fig. 1, in further experiments to avoid the systemic spread of M1 GAS. Mice received a subcutaneous injection of 5×10^5^ KPCY6422c1 cells and were monitored until tumor volumes were within a range of 150-175 mm^3^, around 10 to 12 days, followed by a single intra-tumoral dose of M1 GAS or PBS injection, represented as day 0 post-injection. Tumors were measured every other day for 14 days.

Tumor volume and weight were significantly reduced following treatment with M1 GAS in comparison to PBS control (**Fig. 2A, 2B, 2C**). These results indicate that M1 GAS has a persistent therapeutic effect in more aggressive and clinically relevant subcutaneous KPC-PDAC tumors at medium-size inoculum without systemic spread. Based on this data, we established this dosing to be utilized for further studies.

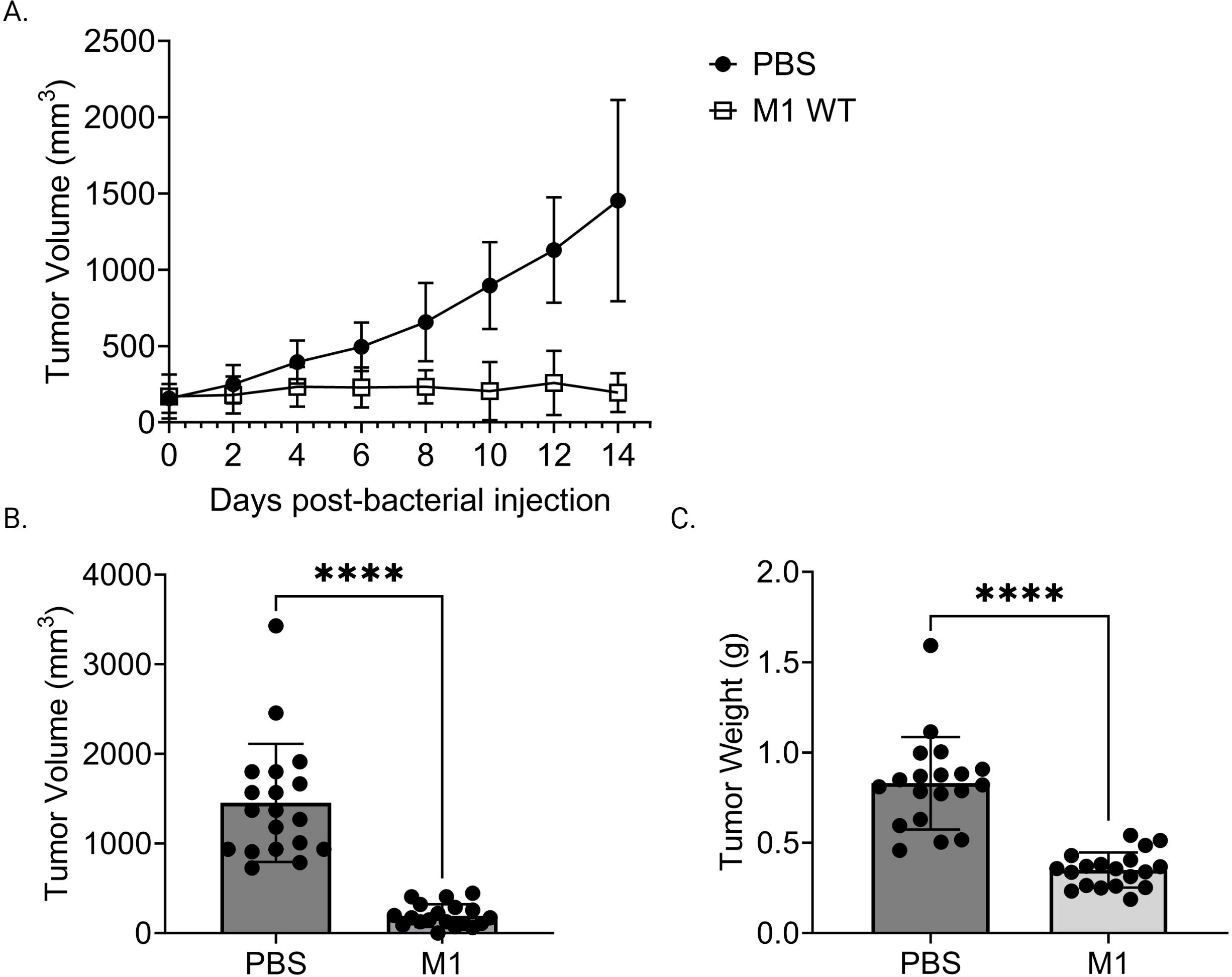

### 3.3 GAS streptococcal collagen-like protein 1 (Scl1) is critical for reduction in PDAC tumor burden and localized tumor colonization

Streptococcal collagen-like protein 1 (Scl1) is expressed by all experimental strains of GAS [41, 42] and is an important adhesion molecule binding to selected type III repeats of cellular fibronectin, a component of the tumoral extracellular matrix also known as oncofetal fibronectin [43, 44]. Given that all three-experimental GAS-M-types used express Scl1, which resulted in tumor regression, we sought to explore if Scl1 was mediating these effects. Following a similar model as above, mice carried subcutaneous KPC-PDAC tumors were administered with a single-dose intra-tumoral injection of either GAS M1 wild-type (WT), an isogenic *Δscl1.1* mutant strain lacking Scl1.1 protein, or a *Δscl1.1::620* mutant strain complemented for Scl1.1 expression. Tumor burden was monitored for fourteen days, then tumors were collected for analyses (**Fig. 3**).

**Figure 3.**
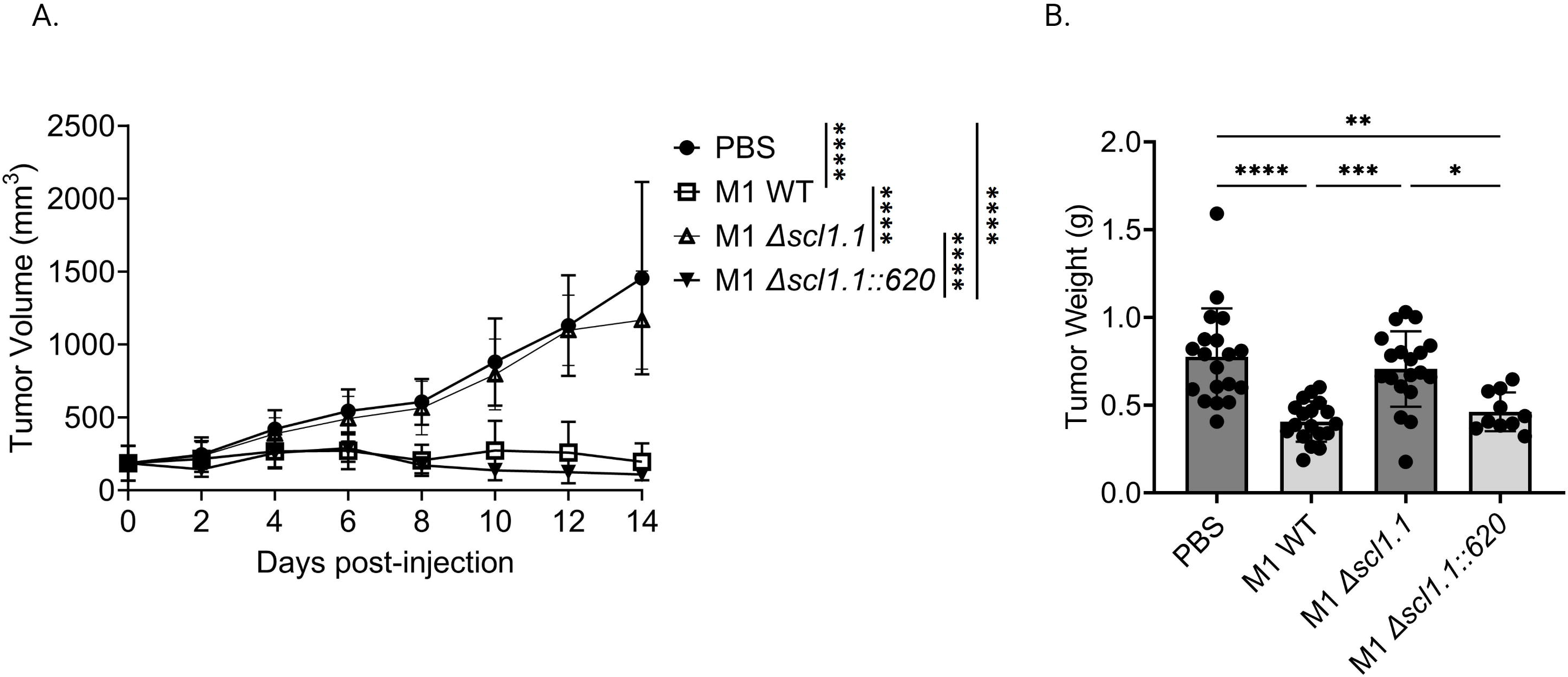
Injection of GAS expressing Scl1 limits the growth of subcutaneous PDAC tumor in mice. (A) Changes in tumor volume following GAS injection. Measurements were taken by caliper during fourteen days following single-dose treatment of indicated GAS strains or PBS control in murine KPC tumors. Results are representative of two independent experiments. Significance is determined by one-way repeated measures ANOVA. ****=p<0.0001. n=10-20 per group total. (B) Tumor weight at experimental endpoint. Mouse tumors treated with indicated strains of GAS or PBS control were extracted and weighed (grams) at day 14. Results are representative of two independent experiments. Each dot represents a single murine tumor. Significance determined by one-way ANOVA with Tukey’s post-hoc test. *=p<0.05, **=p<0.01, ***=p<0.001, ****=p<0.0001

Mice treated with the Scl1-expressing WT bacteria exhibited significant decreases in tumor growth over time as well as decreased tumor weights (**Fig. 3A, 3B**). In contrast, tumor volumes in mice injected with the *Δscl1.1* mutant strain showed no such effect and followed the volume pattern of the PBS control group, suggesting that anti-PDAC activity is contingent on Scl1 expression. When Scl1 expression was complemented back into the mutant strain in mice treated with *Δscl1.1::620*, tumor growth was again diminished and similar to injection with the WT strain, validating a critical role for Scl1 in GAS anti-tumor response.

Based on known Scl1 interactions with tumor extracellular matrix, we hypothesized that Scl1 facilitates a focused nidus of GAS infection, facilitating bacteria localization to the tumor site. Tumor-bearing mice were injected with either GAS M1 WT, an isogenic *Δscl1.1* mutant, or a *Δscl1.1::*620 complemented mutant. Fourteen days post-injection, tumors and spleens were harvested for histopathological analysis and bacterial culture on blood agar (**Fig. 4**). Tumors were locally colonized by GAS strains, as evidenced on Gram-stained histograms and significant recovery of GAS in cultures from tumor homogenates plated on blood agar at the 14-day endpoint post-injection (**Fig. 4A, 4B**). There were no statistically significant differences in tumor colonization levels (CFU per mL) between GAS strains. However, 5 out of 19 (26%) spleen homogenates recovered from mice treated with the isogenic *Δscl1.1* mutant strain yielded bacterial growth, while no growth occurred from the spleens of mice treated with the WT and *Δscl1.1::620* complemented strains (**Fig. C**). These data suggest that Scl1 plays a role in localizing GAS to the tumor microenvironment (TME), thereby limiting the systemic spread of bacteria.

**Figure 4.**
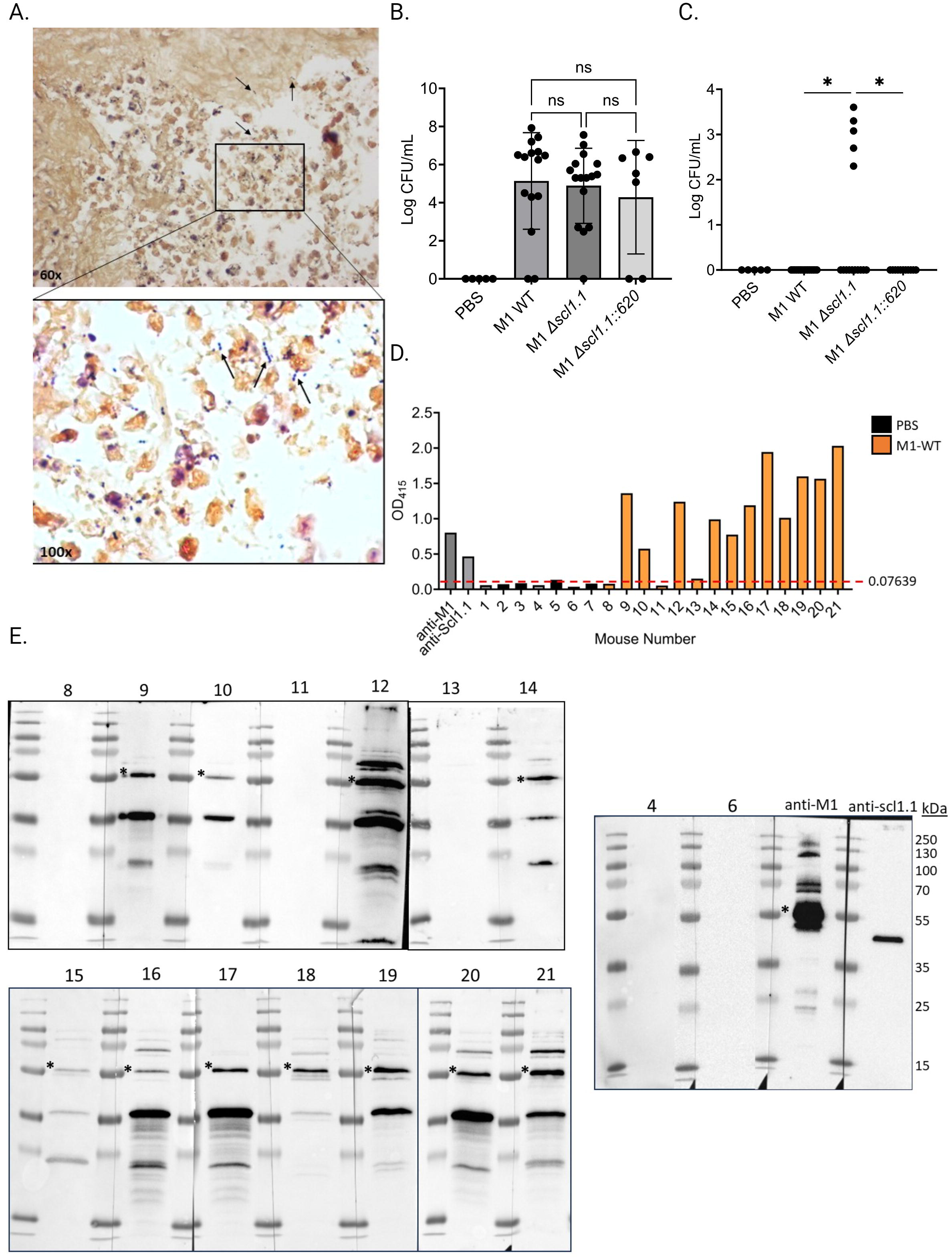
Scl1 promotes localized tumor colonization by GAS without eliciting strong humoral response. (A) Histopathology of GAS-treated KPC tumors. Representative Gram-stained image from the KPC tumor treated with GAS at 60x magnification plus 10x ocular. Arrows indicate GAS at 100x magnification plus 10x ocular under immersion oil. (B) Detection of GAS in KPC tumors. Homogenates were prepared from tumors collected from mice treated with GAS or PBS and serial dilutions were cultured on blood agar, then enumerated. (C) Detection of GAS in spleen homogenates. Spleens were processed and analyzed for GAS presence as above (panel B) for tumor samples. Results are from 2-3 independent experiments. Significance is determined by one-way ANOVA. *=p<0.05. (D) Mouse seropositivity for streptococcal cells developed during tumor colonization by whole-cell ELISA. Mouse sera (1:100 dil.) were collected 14 days after single-dose treatment with either GAS M1 WT (orange bars) or PBS (black bars) and tested for immunodetection of GAS surface antigens; specific anti-M1 (1:1000 dil.) and anti-Scl1.1 (1:1000 dil.) Abs were used as positive controls. Each bar represents the average of three technical replicates. Threshold is indicated by dashed line and was determined by averaging reading of PBS-treated mice at OD _415 nm_. (E) Mouse seroconversion to GAS antigens developed during tumor colonization by western blot analysis. The same mouse sera (1:500 dil.) tested in D were tested against the blots of the M1-GAS cell-wall protein fraction resolved by 15% SDS-PAGE. Asterisk indicates presumed M1-protein immunoreactive band. Specific anti-M1 (1:5000 dil.) and anti-Scl1.1 (1:1000 dil.) Abs were used as positive controls.

Consequently, we next asked whether 14-day-long tumor colonization was sufficient to induce host seroconversion toward M1 bacteria, which would demonstrate immune response to the bacterial injection. Mouse sera (1:100 dil.) collected at the endpoint of the experiment (day 14) following a single-dose injection of GAS or PBS control were screened for anti-M1 GAS immunoreactivity by whole cell ELISA. 11 out of 14 mouse sera (79%) – and none of 7 PBS controls (data not shown) - exhibited seroconversion to whole bacteria (**Fig.4F**). All 14 sera (1:500 dil.) from GAS-treated mice, as well as 7 PBS control sera, were subsequently assessed for antibody reactivity against GAS cell-wall proteins by western immunoblotting (**Fig.4G**). The same 11 mouse sera, previously indicating anti-GAS seropositivity by the whole-cell ELISA, consistently produced three major distinct immunoreactive bands, with one band co-migrating with M1-protein control. As expected, PBS control sera and the remaining 3 sera from GAS-injected mice that were negative by whole-cell ELISA did not show immunoreactive bands against GAS cell wall antigens. Interestingly, the sera analyzed did not show positivity for Scl1.1 antigen in the western blots. Altogether, our data indicate that GAS elicits a host humoral response triggered within immunologically cold TME.

### 3.4 Scl1 is associated with neutrophil infiltration and inhibits neutrophil extracellular trap formation

Given the profound anti-tumor effects of Scl1-mediated GAS bacteria in murine PDAC, we began to consider potential immunostimulatory mechanisms that could be driving this treatment response. We observed that tumors treated with M1 WT and *Δscl1.1::620* complemented strains had higher neutrophil inflammatory indices scored by a blinded pathologist compared to the *Δscl1.1* mutant strain and PBS control (**Fig. 5A, 5B**), indicating more robust neutrophil abundance and suggesting the possibility of fewer NETs (since NETs are a form of neutrophil cell death). Previous studies demonstrated that NETs can enhance pancreatic tumor growth [21] and promote metastasis [18]. We hypothesized that Scl1 may be working through a neutrophil mediated mechanism to exert its anti-PDAC effects. Previously, Scl1 was found to reduce NET formation and killing *in vitro* [24]. Therefore, we sought to further explore the role of Scl1 in cancer-promoting NETs. Bone marrow neutrophils were isolated and incubated with whole cell bacteria (M1 wild-type, an isogenic Scl1-devoid mutant Δ*scl1.1*, and a complemented mutant strain Δ*scl1.1::620*) followed by stimulation with the NET inducer platelet activating factor (PAF). NET production was quantified as a measurement of cell-free DNA (cfDNA) released into the supernatant. Neutrophils co-incubated with the Scl1-expressing bacteria produced significantly less cell-free DNA as compared to the Scl1-lacking mutant and PAF-only control, consistent with reduced NET formation (**Fig. 5C**).

**Figure 5.**
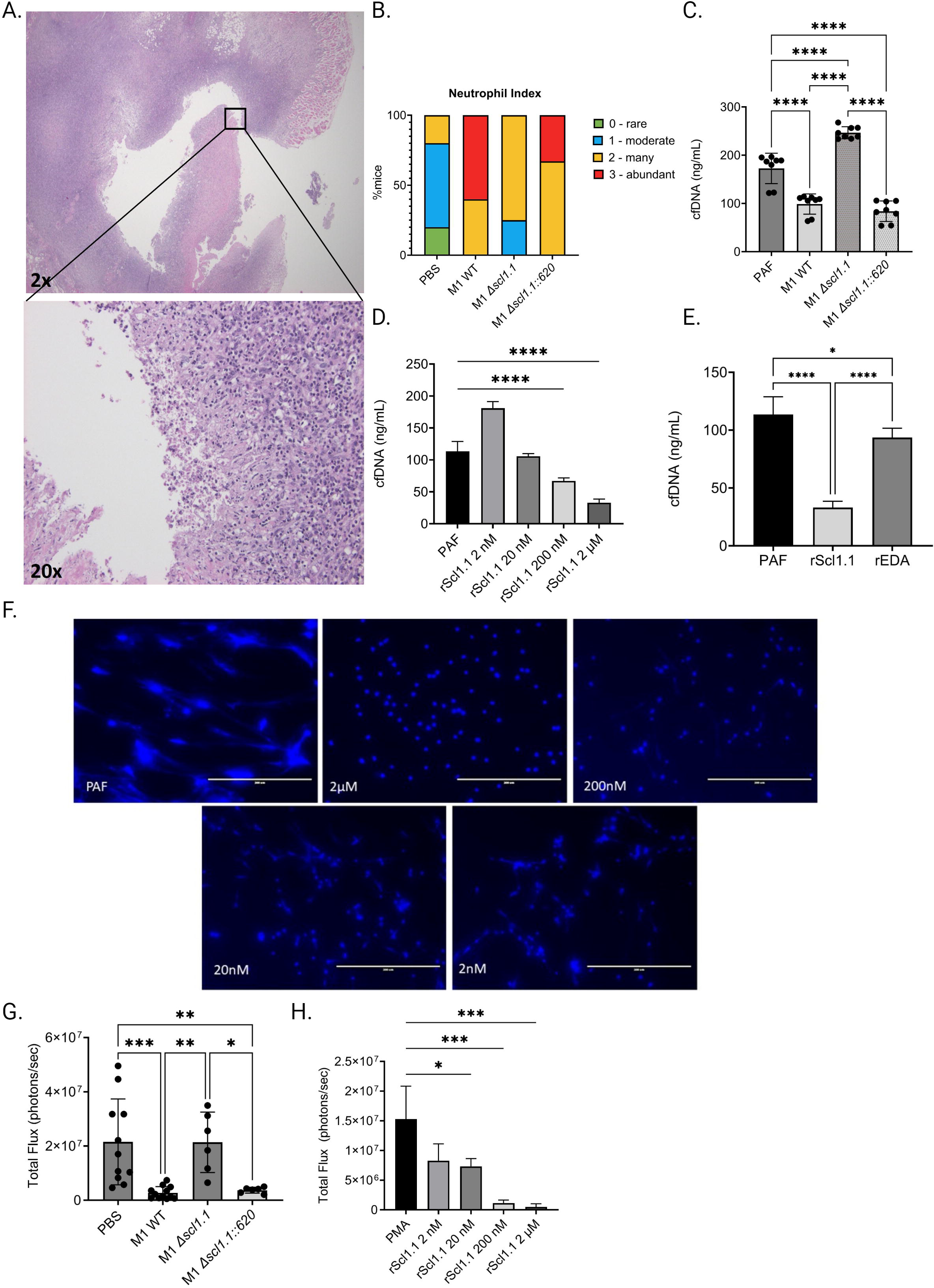
GAS-Scl1.1 impacts intratumoral neutrophil density *in vivo* while rScl1.1 inhibits neutrophil extracellular trap formation *in vitro*. (A) Neutrophil infiltration into M1 GAS treated KPC tumors. Representative H&E stains from tumor specimen at low magnification (top image, 20x) shows neutrophil infiltration at higher magnification (bottom image, 200x). (B) Neutrophil index in KPC tumors. PMN infiltration was scored in tumors from experimental groups harvested 14 days post-injection of a single dose of Scl1 GAS compared to PBS controls (C) NET inhibition induced by *scl1.1*-expressing GAS. *Ex vivo* NET assay measuring cell-free DNA (cfDNA) concentration following co-incubation of neutrophils and designated bacterial strains quantified by Picogreen assay. (D) NET inhibition by rScl1.1. cfDNA concentration following co-incubation of neutrophils and indicated concentrations of rScl1.1. Results are representative of three independent experiments; significance determined by one-way ANOVA with multiple comparison test. ****=p<0.0001. (E) Quantification of NET inhibition by recombinant proteins. cfDNA concentration measured via Picogreen assay of neutrophils co-incubated with rScl1.1 or rEDA followed by stimulation with PAF. Results are representative of two independent experiments (six technical replicates total). Significance determined by one-way ANOVA with multiple comparison test. *=p<0.05, ****=p<0.0001. (F) NET inhibition induced by rScl1.1 *in vitro.* Representative images of neutrophils co-incubated with or without rScl1.1 and stimulated with PAF. NETs are indicated by Hoechst staining. (G) MPO activity of neutrophils isolated from treated, tumor-bearing mice. Total flux as measurement of MPO activity of BMNs isolated from mice 14 days after injection of a single dose GAS treatment or PBS control, demonstrating that Scl1 mediates MPO activity from isolated neutrophils. Results are representative of two independent experiments (n=6 total). Significance is determined by one-way ANOVA and multiple comparison test. *=p<0.05, **=p<0.01, ***=p<0.001 (H) MPO activity of neutrophils treated with rScl1.1. Total flux as measurement of MPO activity of murine BMNs co-incubated with rScl1.1 at indicated concentrations, showing a dose dependent inhibition of MPO. Results are representative of three independent experiments (9 technical replicates total). Significance is determined by one-way ANOVA and multiple comparison test. *=p<0.05, ***=p<0.001.

Because GAS has multiple antigenic influences and mechanisms with the potential to inhibit NET formation [14, 45, 46, 47], we deemed it necessary to confirm Scl1 as the major driver of NET evasion by employing recombinant Scl1 (rScl1) proteins (**Fig. S2**) derived from GAS M1, M28, and M41 strains that inhibited the growth of Panc02 tumors (**Fig. 1**). Bone marrow neutrophils were isolated and incubated with indicated rScl1 proteins. NET release was assessed by measuring the cfDNA and microscopic imaging following fluorescent staining of DNA with Hoechst reagent. Co-incubation of neutrophils with rScl1 proteins significantly inhibited NET formation (**Fig. S3**). rScl1.1 had more potent NET reduction in comparison to other rScl1 proteins tested. Moreover, rScl1.1 inhibited NETs in a concentration-dependent manner (**Fig. 5D, 5F**). In contrast, NET inhibitory activity using unrelated recombinant rEDA protein (**Fig. S2**) as control was not observed (**Fig. 5E**). Overall, we demonstrate for the first time that recombinant Scl1 can inhibit NET formation by the same capability as whole cell bacteria, suggesting strong potential as a useful therapeutic approach for NET inhibition.

We next sought to determine the mechanism of Scl1 NET inhibition. Myeloperoxidase (MPO) activity is a critical step in NETosis whereby MPO stored in the azurophilic granules of neutrophils interacts with neutrophil elastase to induce chromatin decondensation and subsequent NET release [48]. Previous studies demonstrated that Scl1.1 expressed on M1-type GAS suppresses NET formation through inhibition of MPO activity *in vitro* [24]. However, at the current time of publication, these results have not been established in cancer models. Here, we hypothesized that treatment with Scl1-expressing GAS limits MPO activity and subsequent NET formation from neutrophils harvested from our murine model using a novel *ex vivo* bioluminescence assay. Bone marrow neutrophils isolated from mice treated with GAS M1-wild type, the isogenic Scl1-negative mutant and a complemented Scl1-positive strain, or PBS control were stained with the bioluminescence compound luminol, which measures MPO activity [34] and stimulated with phorbol myristate acetate (PMA), a compound known to induce activation of neutrophil NADPH-oxidase and trigger degranulation [49]. Neutrophils isolated from mice treated with the Scl1.1 expressing wild-type and complemented strains exhibited significantly less MPO activity compared to polymorphonuclear neutrophils from PBS controls or treated with the Scl1.1-negative mutant strain (**Fig. 5G**), validating that Scl1 reduces MPO activity.

We next assessed if rScl1.1 protein could elicit similar reductions in MPO activity as seen in our murine model using live bacteria. As described, bone marrow neutrophils were isolated and co-incubated with rScl1.1 at different concentrations followed by luminol staining and PMA stimulation. MPO activity was significantly reduced in neutrophils treated with rScl1.1 in a concentration-dependent manner (**Fig. 5H**). To validate these results, we assessed rScl1.1 in a commercially available MPO inhibitor screening assay, demonstrating that rScl1.1 inhibited MPO production even at low concentrations of protein (**Fig. S4**). Taken together, these results indicate a potential mechanism by which Scl1 inhibits NET formation by hindering MPO activity and offers therapeutic value to rScl1.1.

## 4 Discussion

PDAC is an aggressive cancer characterized by low survival due to failure of early detection and marked treatment resistance. Despite recent advances in cytotoxic regimens, current therapeutic interventions offer minimal treatment response or survival benefit [50], which necessitates exploration into alternative strategies. William Coley established the significance of GAS in treating bone and soft-tissue sarcomas. However, discrepancies in Coley’s work and the development of radiation and chemotherapy brought disfavor towards the use of Coley’s Toxin [7]. In the present work, we elucidate the significance of GAS as a potent anti-cancer agent in PDAC through inhibition of cancer promoting NETosis and re-examine the therapeutic value of bacteriotherapy using GAS.

GAS and other bacteria are attractive candidates for cancer therapy in their capabilities to be recognized by immune cells even in immunosuppressive tumor environments [8]. This recognition has the capacity to switch this immunologically “cold” environment to a more immunostimulatory one [51]. As demonstrated by Maletzki *et al.* (2008) using Panc02 subcutaneous model, a single intra-tumoral injection of M49-type GAS can mediate anti-tumor effects through direct tumor lysis and stimulation of anti-tumor immunity by active infection. In the present study, injection with M1, M28, and M41 GAS each significantly reduced Panc02 tumor growth. Moreover, a subsequent use of the M1 GAS led to tumor reduction of a more aggressive and clinically relevant cell lineage such as the KPCY6422c1 line, which solidifies this concept and offers greater potential for clinical translation.

A hallmark of PDAC is an immunosuppressive tumor micro-environment that renders tumors resistant to current immunotherapies [52, 53]. Existing literature demonstrates the contribution of NETs to this immunologically “cold” state [54, 55] and their enhancement to cancer progression [17, 56, 57] and reduced patient survival [23, 58]. Thus, NETs are encouraging targets for further cancer therapies. The current work strengthens existing data generated in a non-cancer inflammatory model to suggest that Scl1 on GAS cells is a potent NET inhibitor through reduction in MPO activity [14, 24]. Based on these findings, we tested a recombinant Scl1 protein (rScl1.1) for potential NET inhibition. Neutrophils co-incubated with rScl1.1 had a decreased propensity to form NETs indicated by reduced cell-free DNA concentrations similar to that from Scl1-expressing live bacteria. Moreover, rScl1.1 inhibited MPO activity indicated by our *in vitro* luminol assay. Taken together these results suggest that rScl1.1 functions in the same manner as the native protein expressed in live bacteria. This original finding potentiates rScl1.1 as a conceivable PDAC therapeutic agent to combat concerns accompanying live and whole bacterial therapy and represents an innovative advance in the field, as recombinant Scl1 has not been previously tested to inhibit cancer-associated NET formation.

Our previous work reported that Scl1 selectively binds to cancer-associated fibroblast (CAF)-deposited oncofetal fibronectin, [44, 59, 60] which can modulate the tumor extracellular matrix microenvironment [43]. Here, we observed that injection of Scl1-expressing GAS was associated with tumor-localized nidus of infection persisting for an extended time, whereas Scl1-lacking isogenic mutant had greater systemic bacterial dissemination. Despite these encouraging results in preclinical models in the current study, injection of live GAS may not be a translatable long-term strategy in humans given the potential for pathogenic infection. Our findings implicated Scl1 in the therapeutic efficacy of GAS, with Scl1-devoid mutant strain demonstrating no significant treatment response comparable to PBS control. Taken together with the evidence demonstrating robust neutrophil infiltration at sites of tumor colonization by GAS, these results potentiate the capacity of Scl1 to counteract the immunotolerant environment and function as an anti-cancer therapeutic. Further analysis elucidating specific immune infiltrates and inflammatory responses in subsequent studies will offer a more mechanistic approach to characterizing immunomodulatory responses within the tumor microenvironment in response to GAS bacteriotherapy.

Of interest when considering future therapeutic strategies, we observed host immunity to GAS antigens, thus, increasing the likelihood of resolution of inflammation and the need for additional treatment administration to induce a durable response. Repeat treatment within 3 weeks is well in line with currently utilized cytotoxic treatment regimens and is clinically realistic. Experimental treatments utilizing other bacteria, such as *Listeria* [61], show promise and require multiple rounds of injection as well. However, this observation expresses a need to understand specific host-immune responses to bacterial antigens to identify proper dosing. Herein, we report that mice seroconvert to live bacteria. While this finding is not surprising given the prominent antigenicity of M proteins [32, 62], it proposes potential challenges to future studies examining the efficacy of GAS therapy. However, highly immunogenic antigens are beneficial in the context of vaccines [63]. Therefore, we cannot exclude the possibility of utilizing M-proteins in cancer vaccine development to elicit potent antitumor immunity. Surprisingly, western blot analysis did not demonstrate seroconversion to Scl1.1 in this sample set. Although the anti-tumor effects demonstrated herein are suggested to be Scl1-mediated, identifying the complex immunological response within the tumor microenvironment is necessary to determine immunotolerance at later intervals of treatment. Given the lack of seroconversion demonstrated in these early studies, future studies investigating the therapeutic efficacy of Scl1.1 can be conducted utilizing rScl.1 or engineered non-pathogenic or attenuated bacteria expressing Scl1.1, as opposed to live M1-WT GAS.

In conclusion, we demonstrate a profound anti-tumor effect of several GAS strains, validated in two PDAC model cell lines. We also identify a novel role for Scl1.1 in PDAC therapy and provide insight on future applications utilizing rScl1.1 to target cancer-promoting NETs. We also address limitations of current and future bacteriotherapies through analysis of seropositivity.

Taken together, Scl1.1 could be a beneficial therapeutic approach in PDAC that warrants additional studies.

## 7 Acknowledgments

We acknowledge James (Jim) Dale, Division of Infectious Diseases at the University of Tennessee Health Science Center for the anti-M protein 30-valent antibodies used as controls in the ELISA and western blots. We also thank all additional members of the Liu, Lukomski, and Boone labs for their assistance in experiments. Some figures were configured using Biorender.

## 9 Data Availability Statement

The original contributions presented in the study are included in the article/supplementary material, further inquiries can be directed to the corresponding authors.

