## Supplemental Figures for "Group A Streptococcal Collagen-like Protein 1 Restricts Tumor Growth in Murine Pancreatic Adenocarcinoma and Inhibits Cancer-Promoting Neutrophil Extracellular Traps"

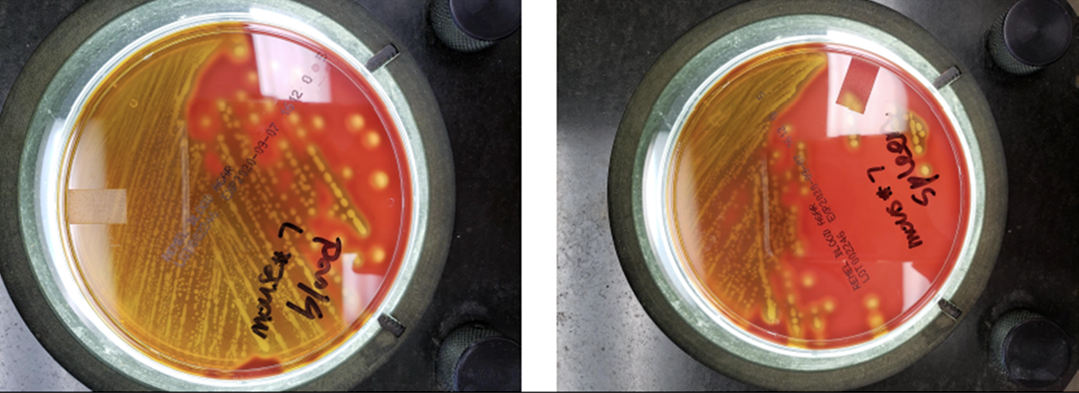

**Supplementary Figure 1. Detections of M1 GAS in blood and organs.** Images of blood and spleen homogenate from representative infected mouse were cultured on blood agar yielding β-hemolytic colonies confirming GAS presence in mouse specimen 7.

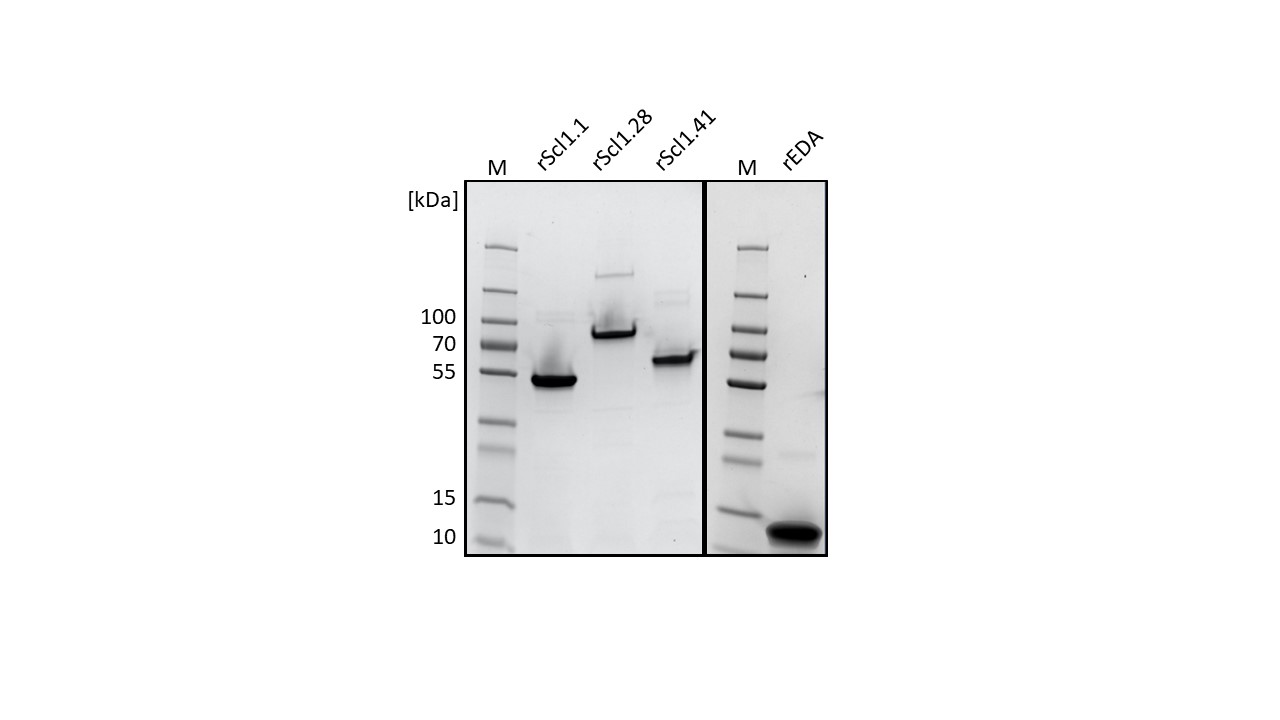

**Supplementary Figure 2. Purified recombinant proteins.** 4-20% SDS-PAGE confirming purity and integrity of rScl1 and rEDA preparations. rScl1.1 corresponds to Scl1 in M1-type strain, rScl1.28 in M28-type strain, rScl1.41 in M41-type strain. M, molecular mass standard in kDa.

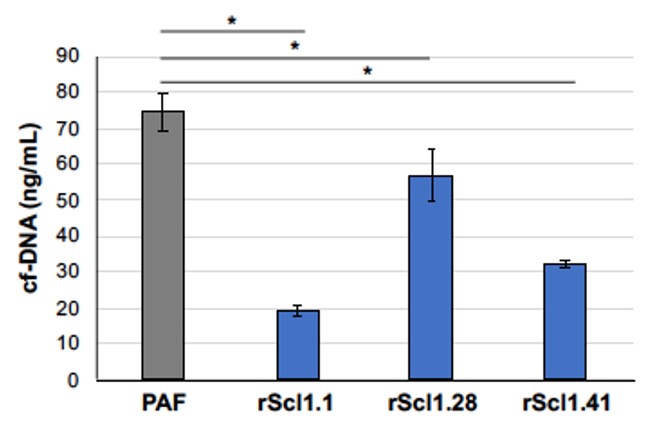

**Supplementary Figure 3. Recombinant Scl1 proteins inhibit NET formation.** Effect of rScl1 proteins on NET formation as a quantification cfDNA concentration by Picogreen assay following co-incubation of neutrophils isolated from bone marrow of mice and indicated rScl1 proteins. Significance determined by one-way ANOVA with multiple comparison test. *=p<0.05

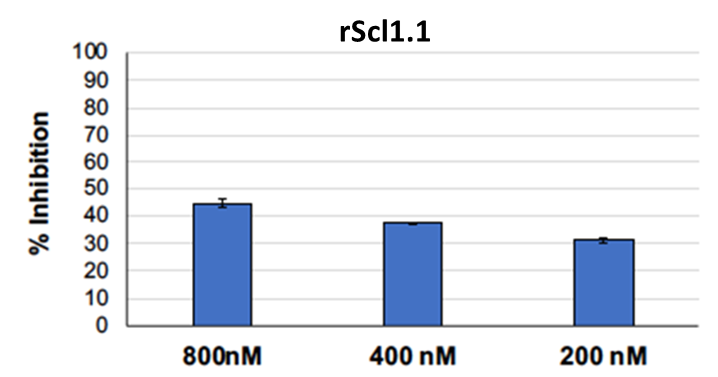

**Supplementary Figure 4. Recombinant Scl1.1 inhibits MPO activity.** Effect of rScl1.1 on MPO activity as a percentage of MPO inhibition using a commercially available biochemical screening assay.
